# A chromosome-level genome assembly of *Thecaphora frezzii*, cause of peanut smut, reveals the largest genome among the true smut fungi

**DOI:** 10.64898/2026.02.02.703329

**Authors:** Nicholas Greatens, M. Brian Couger, Mariano Maestro, Guillermo Cabrera Walsh, Sergio Morichetti, Luke J. Tallon, Rebecca S. Bennett, Josh P. Clevenger, Kelly D. Chamberlin, Rachel A. Koch Bach

**Affiliations:** Foreign Disease-Weed Science Research Unit, USDA Agricultural Research Service, Fort Detrick,MD, U.S.A.; Oak Ridge Institute for Science and Education, ARS Research Participation Program, Oak Ridge,TN, U.S.A.; Department of Thoracic Surgery, Brigham & Women’s Hospital, Boston, MA, U.S.A.; Foundation for the Study of Invasive Species, Hurlingham, Buenos Aires Province, Argentina; Sanchez Farms, Old Town, FL, U.S.A.; Institute for Genome Sciences, University of Maryland, Baltimore, Maryland, U.S.A.; Peanut and Small Grains Research Unit, Oklahoma & Central Plains Agricultural Research Center, USDA Agricultural Research Service, Stillwater, OK, U.S.A.; HudsonAlpha Institute for Biotechnology, Huntsville, AL, U.S.A.

**Keywords:** Effectors, comparative genomics, Transposable elements, *Ustilago*, *Pseudozyma*, phylogenomics

## Abstract

Peanut smut, caused by the fungus *Thecaphora frezzii*, is a significant disease of peanuts in Argentina. Infected plants have seeds replaced by a mass of dark teliospores, reducing yield and seed quality. To prevent the spread of the pathogen, several countries have limited import of raw peanuts from Argentina, a major grower and exporter. Following successful *in vitro* culture of the fungus in its haploid stage, we produced a chromosome-level genome assembly of the species for the first time. We compare this genome with those of 49 other species of true smut fungi, or Ustilaginomycetes, including species of medical, agricultural, and industrial importance, some of which are known as pathogens and others only as saprotrophic yeasts. At almost 39 Mb, *T. frezzii* has the largest genome of the smut fungi sequenced to date and the highest repetitive content. While it shares some core effectors with species of the distantly related and better studied *Ustilago* and related fungi, predicted effectors only found in *T. frezzii* or in *Thecaphora* suggest unique infection strategies. Comparisons among the 50 smut genomes also show that the 14 smut fungi observed only as yeasts share genomic traits such as low repeat content and generally smaller genomes. This supports the hypothesis that some smut fungi are adapted to saprotrophic growth as yeasts. The high-quality, annotated genome for *T. frezzii* will be a valuable resource for investigating the population dynamics and evolution of an economically important pathogen, as well as illuminating an understudied clade of smut fungi.

**Article summary:** Peanut smut is a destructive and costly disease of peanuts in Argentina. For the first time, a high quality, annotated genome is presented for the causal agent, *Thecaphora frezzii*. This fungus has the largest and most repetitive genome of the true smut fungi, thus prompting comparison with 49 other species of smut fungi with available genomes, including non-pathogenic ones. While it shares some likely pathogenicity factors with well-studied smuts, it has many unique genes, a trait reflective of its evolutionary distance and likely novel mechanism for infection. This important genomic resource will benefit research regarding the evolution and adaption of *T. frezzii*, the development of diagnostic tools that enable rapid detection of it, and the study of smut fungi broadly.

## Introduction

Ustilaginomycetes, or the true smut fungi, is a large and important class of plant pathogens. Most species fall within two large orders, the Ustilaginales and the Urocystidales. The Ustilaginales includes notable smut species like *Mycosarcoma maydis* (formerly *Ustilago maydis*), an important model species and cause of corn smut, and *Sporisorium scitamineum*,cause of sugarcane smut. The Urocystidales includes early diverging species like *Thecaphora solani*, the cause of potato smut. Smut fungi typically have a complex, dimorphic life cycle. One phase is haploid, where the fungus is saprotrophic and generally yeast-like. While some Ustilaginomycetes have only been observed in this stage, most described species also have a dikaryotic pathogenic phase that is initiated after fertilization and results in the production of melanized, powdery teliospores adapted for dispersal and survival. As plant pathogens, smut fungi are biotrophic, meaning they do not kill their hosts outright, but they may still cause significant damage and reductions in yield.

One smut fungus species, *Thecaphora frezzii*, within the Urocystidales, is an emerging threat to peanut production. *Thecaphora frezzii* causes peanut smut, a major disease of peanut (*Arachis hypogaea*) in Argentina (Rago et al. 2017; Ospina-Maldonado et al. 2022; Paredes et al. 2024). It was first found in 1959 in southern Brazil in the pods of an endemic wild peanut species, *Arachis kuhlmannii* (Krapovickas et al. 2005), but the fungus was not observed in commercial peanut fields until the fall of 1995 (Marinelli et al. 1995; Rago et al. 2017; Paredes et al. 2024). Since then, *T. frezzii* has spread from a small area in the province of Córdoba to all commercial peanut production areas across multiple provinces, developing into a serious and costly problem for the Argentine peanut industry (Marinelli et al. 2010; Rago et al. 2017; Cazón et al. 2018). In 2024, Argentina ranked seventh in production and second in exports for peanuts worldwide (USDA Foreign Agricultural Service, 2024). Because of the risk posed by peanut smut disease, several countries, including Australia, South Africa, and the United States, have imposed import restrictions on raw or unprocessed peanuts from Argentina, along with Brazil and Bolivia, where *T. frezzii* was also observed on wild peanut species (Arias et al. 2021; Australian Biosecurity Import Conditions 2017; Oilseeds Advisory Committee 2022). Only Arachis spp. are known to be infected by *T. frezzii* (Paredes et al. 2024).

Infection is believed to occur after flower production when the peanut plant produces a peg, or gynophore, that penetrates infested soil (Marinelli et al. 2008). Teliospores germinate upon recognition of peanut pegs and produce haploid filamentous hyphae that fuse with those of a compatible mating type, undergoing plasmogamy. The dikaryotic hyphae infect the peg and grow within the host as the fruit develops (Arias et al. 2021; Paredes et al. 2024). In infected plants, peanut seeds within the pods are partially or completely replaced by a mass of teliospores (Arias et al. 2021; Cazón et al. 2018). Infected pods are sometimes but not always hypertrophic (Bennett et al. 2021), and aboveground plant parts do not show any discernable symptoms, a factor which may have enabled the disease to spread undetected (Paredes et al. 2024).

While many smut fungi are amenable to culturing, especially in the haploid stage of the life cycle, *T. frezzii* is not easily cultured using typical methods. This trait has hindered the extraction of sufficient high-molecular weight DNA needed for producing a high-quality, contiguous genome. Some scientists, however, reported success inducing teliospore germination using media augmented with plant extracts (Arias et al. 2021; Rago et al. 2017; sources within). From a stable, haploid culture generated from a teliospore germinated in vitro and propagated in liquid media, we produced and annotated a chromosome-level genome assembly of *T. frezzii* for the first time. The annotated genome will be a valuable resource to study peanut smut disease as well as the understudied Urocystidales.

## Materials and Methods

### Culturing and extraction

*T. frezzii* isolate ‘Nelio’ was collected from a peanut pod grown in the 2020-21 season from a highly infested field near Rio Cuarto, Córdoba, Argentina (-33.1071, -63.9757). The pod was surface sterilized in 10% NaOCl for five minutes and opened by hand. Teliospores were collected into cryovials and stored at 4°C. A peanut peg extract was prepared by rinsing young peanut pegs with water, crushing with a mortar and pestle, incubating at 40°C for 2 hours in sterile distilled water, and sterilizing the peg solution with a 0.22 nm syringe filter. The extract was spread on the surface of potato dextrose agar (PDA) containing 20 g/L sucrose and 0.5 g/L chloramphenicol. Teliospores were surface sterilized in 10% NaOCL for 5 min, rinsed three times in sterile distilled water, and spread onto plates with a Drigalski spatula. After two weeks of incubation at 25°C, a single germinated teliospore was transferred to and maintained on PDA through subculturing.

Seven 100 mL flasks of potato dextrose broth (PDB) were inoculated with macerated *T. frezzii* mycelium from the pure, haploid isolate to generate enough tissue for DNA and RNA extractions. Cultures were maintained in dark on a shaker for 12 days at ambient temperature, after which the tissue was collected in an EASYstrainer (Greiner Bio-One, Monroe, North Carolina) and ground immediately in liquid nitrogen. DNA was extracted with a cetyltrimethylammonium bromide (CTAB) protocol (Gardes and Bruns 1993) that included an RNAse treatment. Aliquots of genomic DNA were pooled together and purified with the Monarch Nucleic Acid Purification Kit (New England BioLabs, Ipswich, Massachusetts). Fragments < 500 bp were removed with AMPure PB beads (Pacific Biosciences, Menlo Park, California). The library was prepared with the SMRTbell Prep Kit 3.0 per manufacturer instructions, and size-selected with AMPure PB beads to retain fragments > 5000 bp. The DNA library was sequenced on a PacBio Revio. RNA was extracted with the E.Z.N.A. Fungal RNA Mini Kit (Omega Bio-Tek, Norcross, Georgia) and sequenced on the Illumina NovaSeq 6000 with an S4 flow cell (Illumina, inc. San Diego, CA). PacBio and RNA sequencing were conducted at Maryland Genomics (part of the University of Maryland School of Medicine Institute for Genome Sciences). Hi-C reads were generated by Phase Genomics (Seattle, Washington) using an Illumina NovaSeq X Plus 10B. Raw reads were deposited in the Sequencing Read Archive (SRA) of the National Center for Biotechnology Information (NCBI) under BioProject PRJNA1250501.

### Genome assembly and annotation

PacBio HiFi genomic reads were trimmed of adapters and filtered for > Q15 by the sequencing core at Maryland Genomics. Quality was examined with NanoPlot v1.42.0 (De Coster and Rademakers 2023). Reads were assembled with Hifiasm v0.18.9 with default settings (Cheng et al. 2021). The primary assembly was assessed with blobtoolkit v4.3.5 (Challis et al. 2020) using Diamond (Buchfink et al. 2021) with the UniProt database (The UniProt Consortium et al. 2023), BLASTn (Camacho et al. 2009) with the NCBI nucleotide library, and minimap2 (Li 2018) with recommended parameters. After inspection of BLASTn results and the GC-coverage plot, scaffolds were filtered for > 10x coverage and > 45% GC content. Contigs below the 45% threshold were derived from the mitochondrial genome. No contaminants were detected within the assembly. Hi-C reads were mapped to the contigs using the pipeline developed by Phase Genomics (https://phasegenomics.github.io/2019/09/19/hic-alignment-and-qc.html). Contigs were scaffolded with YAHS using default settings (Zhou et al. 2023), and scaffolds were inspected and manually curated in Juicebox (Durand et al. 2016) based on Hi-C contacts. Assembly quality was assessed with compleasm v0.2.6 (Huang and Li 2023) using the basidiomycota_odb10 dataset. The FindTelomeres.py script (Sperschneider, https://github.com/JanaSperschneider/FindTelomeres) was used to detect telomeres.

Repetitive elements were annotated with EDTA (Ou et al. 2019) with the sensitive setting, employing RepeatModeler (Flynn et al. 2020), LTRFinder (Xu and Wang 2007), LTRretriever (Ou and Jiang 2018), Generic Repeat Finder (Shi and Liang 2019), TIR Learner (Su et al. 2019), HelitronScanner (Xiong et al. 2014), and TESorter (Zhang et al. 2022). The assembly was soft-masked with EDTA/utils/make_masked.pl. Genes were predicted using the Funannotate v1.8.16 pipeline (Palmer and Stajich 2023) with the train, predict, and update modules using default settings with the RNA-Seq data as additional input. Funannotate uses Trinity (Grabherr et al. 2011) and StringTie (Pertea et al. 2015) for transcript assembly, and Augustus (Stanke et al. 2006), CodingQuarry (Testa et al. 2015), and Pasa with EvidenceModeler (Haas et al. 2008) for gene prediction. The longest isoforms were retained (AGAT: agat_sp_keep_longest_isoform.pl; Dainat 2022). Functional annotation was added using data from eggNOG-mapper v2 (Cantalapiedra et al. 2021) and the eggNOG 5.0 database (Huerta-Cepas et al. 2019), interproscan v5.67-99 (Jones et al. 2014), antiSMASH v7.1.0 (Blin et al. 2023), and SignalP (Teufel et al. 2022). The annotated nuclear genome was deposited in NCBI GenBank as accession JBPEIW000000000.

Carbohydrate-Active Enzymes, or CAZymes, were also predicted independently from the Funannotate pipeline using dbCAN v4.1.4 (Zheng et al. 2023), with genes supported by two of the three methods retained as candidates. Family level annotations were taken from the best supported prediction by dbCAN, or else from HMMER hits if the dbCAN classification was not available. Effectors were predicted with EffectorP v3 (Sperschneider and Dodds 2022) from predicted secreted proteins. Additionally, secreted proteins orthologous to novel secreted core effectors in the Ustilaginaceae, or the family containing *Ustilago* and *Mycosarcoma* maydis, previously characterized by Schuster et al. (2024), were considered candidate effectors. The proximity of genes encoding predicted effectors and BUSCO (Simão et al. 2015) genes to repetitive elements was assessed using bedtools slop and bedtools intersect with the EDTA repeat annotation files *EDTA.TEanno.gff3 and *EDTA.intact.gff3.

### Comparative genomics

Fifty genomes of Ustilaginomycetes designated by NCBI as reference genomes as of June 3, 2024 and genomes from three additional species hosted by the Joint Genomes Institute were downloaded and used for initial comparative genomics analyses. Genomes were assessed for quality and completeness with gfastats (Formenti et al. 2022) and compleasm (Huang and Li 2023) using the basidiomycota_odb10 BUSCO dataset. Of these 53 genomes, four were excluded from later analyses: *Fereydounia khargensis* and *Ustilago trichophora* were excluded due to irregular BUSCO scores (low completion (55%) and high duplication (88%), respectively); *Moesziomyces* sp. 2 was excluded since submitters indicated it was likely the same species as *Moesziomyces* sp. 1; and *Urocystis occulta*, was excluded after the genome appeared unexpectedly within the Ustilaginales in preliminary phylogenetics analyses, a position inconsistent with phylogenetic data derived from the type specimen for the species. Our genome was used for *T. frezzii*. In total, 50 Ustilaginomycetes reference genomes representing 50 species were included in the analyses (Table S1).

The BUSCO phylogenomics pipeline (https://github.com/jamiemcg/BUSCO_phylogenomics) was used with the 50 Ustilaginomycetes genomes and the outgroup, *Malassezia globosa*, in the *Malasseziomycetes*. With the unrooted gene trees for the 1764 Basidiomycota BUSCO genes as input, ASTRAL-IV v1.22.4.6 (Zhang and Mirarab 2022) with CASTLES-2 (Tabatabaee et al. 2023) was used to infer a phylogenetic tree with a multi-species coalescent model.

While some reference genomes had annotations available, all genomes were independently annotated using the same pipeline for consistency. Soft-masking was removed and repeat content was assessed with EDTA with the sensitive setting, as above. Assemblies were soft-masked with the EDTA repeat libraries and submitted to BRAKER v3.0.3 (Gabriel et al. 2024) with the basidiomycota_odb11 dataset as protein hints and with Augustus trained on *M. maydis*. Features of GFF3 files output by Braker were summarized with AGAT agat_sq_stat_basic.pl (Dainat 2022), after filtering for the longest isoforms. Predicted proteomes were submitted to SignalP 6.0 to predict secreted proteins (Teufel et al. 2022). From the predicted secreted proteins, effectors were predicted using EffectorP 3.0 (Sperschneider and Dodds 2022) and CAZymes were predicted with dbCAN v4.1.4 as described above. The genes encoding secreted proteins and predicted apoplastic and cytoplasmic effectors were summarized for each species. Gfastats, compleasm, AGAT, EDTA, dbCAN, and EffectorP outputs for all genomes were summarized and aggregated with custom python and shell scripts. Species were designated as “plant pathogenic”, “no plant pathogenicity observed”, or “no data” based on literature searches or from data uploaded to NCBI with genome submissions (Table S1).

To assess shared effector-encoding genes and other genes of interest, orthofinder (Emms and Kelly 2019) was run with the *M. maydis* RefSeq proteome (GCF_000328475.2), the predicted *Thecaphora thlaspeos* proteome (personal communication, Vera Göhre), the *T. frezzii* predicted proteome from the Funannotate pipeline, and the 47 BRAKER3 annotated genomes.

All scripts and the data necessary to produce figures are available in github.com/ngreatens/Thecaphora_frezzii_genome or at doi://10.5281/zenodo.15677049.

## Results

### *Thecaphora frezzii* genome

The genome was assembled into 40 contigs and 29 scaffolds, totaling 38.79 Mb, the largest sequenced smut genome at the time of writing (Fig. 1; Table S2). Nineteen scaffolds are assembled end to end, representing chromosomes, while eight scaffolds lack one telomere (Fig 1.). A hi-C contact map shows the 27 largest scaffolds (with Fig S1) are highly contiguous without few links between scaffolds, but the absence of terminal telomeric sequences on some scaffolds precludes a definitive conclusion on the number of chromosomes. Scaffolds range in size from 415 kb – 2.6 Mb. The haploid genome assembly is complete or nearly complete, with 99.32% of Basidiomycota BUSCO genes complete and present in the assembly. Of these, 96.2% are single copy, 3.12% are duplicated, 0.4% are fragmented, and 0.28% are missing (Table S3).

**Figure 1.**
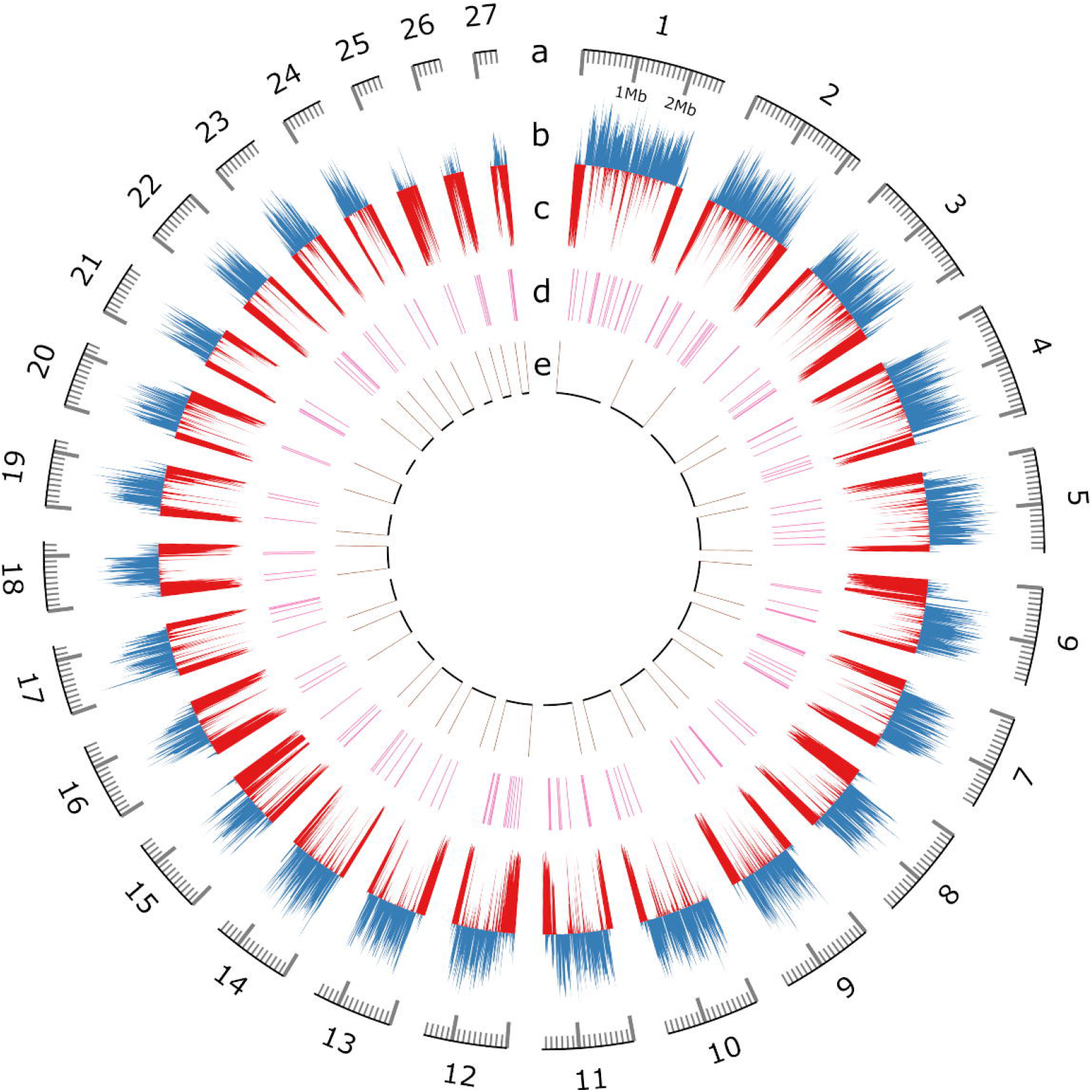
Circos plot of the *Thecaphora frezzii* reference genome, showing: (a) scaffold lengths; (b) percent coding DNA sequences in 10kb windows; (c) percent transposable elements in 10kb windows (reversed); (d) locations of candidate effector-encoding genes; and (e) telomeric sequences on scaffold ends. The plot was prepared with the python package pyCirclize (https://github.com/moshi4/pyCirclize).

The *T. frezzii* genome has significant repeat content, with 12.5 Mb of repeat elements, representing 32.2% of the genome (Figs 2,3; Table S4). Long terminal repeat retrotransposons (LTRs) are the most highly represented among the repetitive elements, making up 13.8% of the genome, with Ty1/copia LTRs being the most abundant type (Table S4). Forty-one percent of the LTRs, representing 5.7% of the genome, could not be classified to the superfamily level, but were recognized as LTRs. Repeat fragments, or sequences homologous to other repetitive elements but structurally incomplete, make up 10% of the genome.

**Figure 2.**
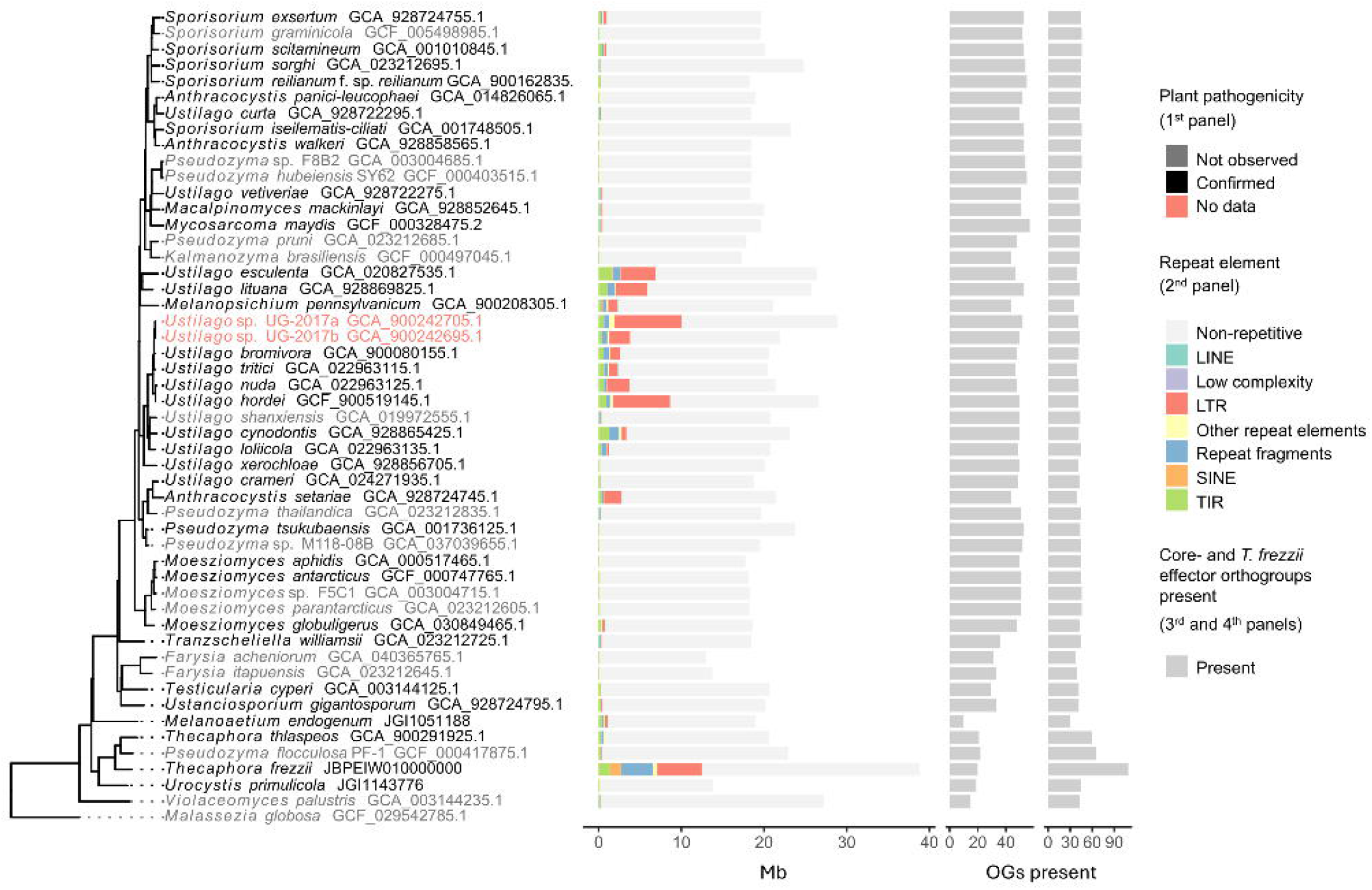
Phylogenetic tree of 50 Ustilaginomycetes with quantities and classes of repeat elements (panel 2), and presence of Ustilaginaceae core- and *T. frezzii* effector orthogroups (OGs; panels 3 and 4). The phylogenetic tree was generated using a multispecies coalescent model (Astral-IV) for the 50 Ustilaginomycetes species based on gene trees for 1122 fungi BUSCO genes (github.com/jamiemcg/BUSCO_phylogenomics). The tree is rooted on *Malassezia globosa*. Species names are black, gray, or red to indicate observed plant pathogenicity. Species names appear as given in NCBI, as of June 4, 2025.

**Figure 3.**
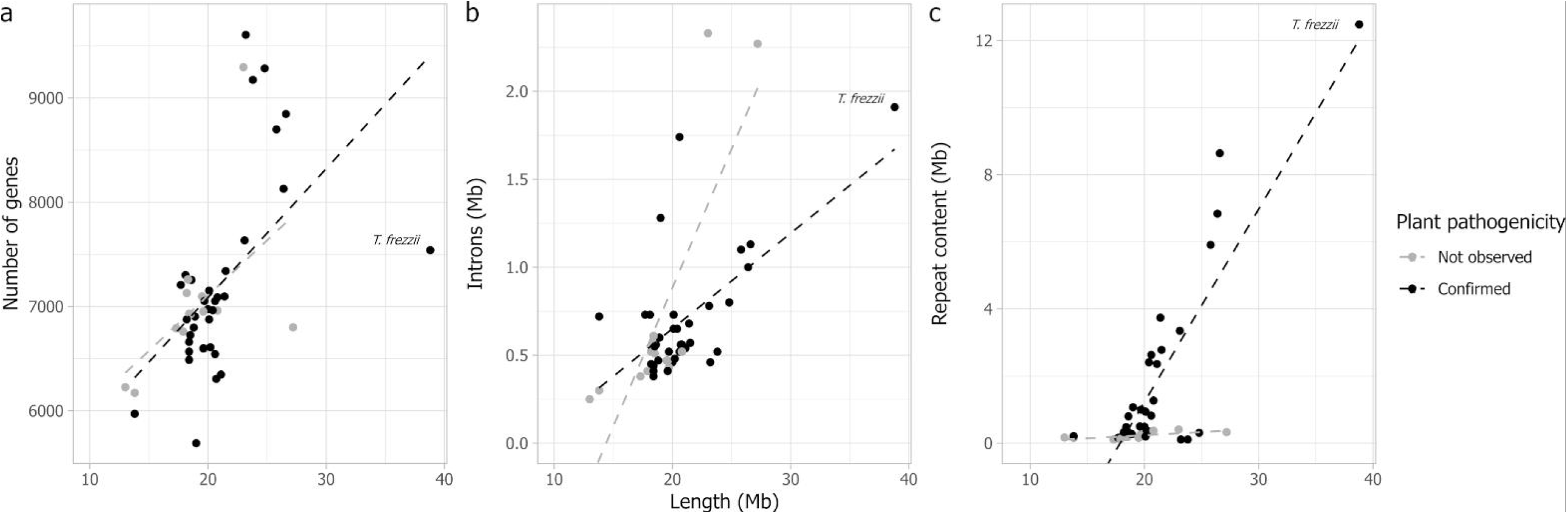
Scatter plots that compare genomic features with genome size for 50 true smut fungi. Species are colored based on their observed plant pathogenicity. Gene and intron counts are based on BRAKER-3 annotated genomes. (a) Number of predicted genes vs genome size (b) Introns (Mb) vs. genome size (c) Repeat content (Mb) vs genome size.

AntiSMASH predicted ten biosynthetic gene clusters encoding secondary metabolites: one fungal RiPP (Ribosomally synthesized and post-translationally modified peptide), one polyketide, two terpenoids, and six non ribosomal peptides. A biosynthetic gene cluster orthologous to those producing ustilagic acid in *M. maydis* and flocculosin in *Pseudozyma flocculosa* is conserved in *T. frezzii*, with the core biosynthetic fatty acid synthase gene most closely related to the fas2 ortholog in *P. flocculosa*. Clusters that synthesize mannosylerythritol lipid (MEL) and itaconic acid synthesis in *M. maydis* are not conserved in *T. frezzii*, although orthologs for *UMAG_03115*, a major facilitator involved in MEL synthesis, and *UMAG_05074* (*cyp3*), a core biosynthetic gene in itaconic acid biosynthesis, are present: *ACQY0O_001991* and *ACQY0O_000524*, respectively. One hundred and ninety-three CAZymes were predicted, of which 75 had a secretion signal, and eight were predicted to be apoplastic effectors (Table S5). Mating-type (MAT) loci were identified and are on separate chromosomes, indicating a tetrapolar mating system.

Five hundred and fourteen genes in *T. frezzii* were predicted to encode secreted proteins. Of these predicted secreted proteins, 150 were predicted as effectors by EffectorP (Table S6). Thirty-six were predicted to be localized to the apoplast and 82 to the cytoplasm, while 32 were predicted as either Apoplastic/cytoplasmic or Cytoplasmic/apoplastic effectors, indicating some uncertainty. In *M. maydis*, 73 genes in 53 orthogroups were identified by Schuster et al. (2024) as core effectors of the Ustilaginaceae based on secretion signal and absence of known protein domains. In our separate orthofinder analysis using the predicted proteomes for the 50 Ustilaginomycetes, the same 73 genes were inferred to be in 57 orthogroups (Table S7). Twenty-seven proteins in *T. frezzii* are orthologous with core effectors in *M. maydis* and were also considered candidate effectors if they contained secretion signals, regardless of EffectorP predictions (Table S7-8). Six of these 27 genes encode orthologs of the conserved effectors pep1, sta1, and stp1 (cce1), stp3, and stp4, although orthologs for pep1 and sta1 are not predicted as secreted. Three others are orthologous to *UMAG_12316* or *UMAG_12205*, which contribute to virulence in *M. maydis* (Schuster et al. 2024), but are not predicted as secreted. Only five proteins predicted as effectors by EffectorP were orthologous with core effectors in *M. maydis* (Table S7). In total, using the two methods, 165 candidate effectors were identified in 120 orthogroups.

An ortholog of PFL1_02826, an effector produced by *P. flocculosa* Pf-1 that is required for biocontrol of powdery mildew (Santhanam et al. 2023), is present in *T. frezzii* (ACQY0O_008349) and has a secretion signal but was not predicted as an effector. Orthologs of Pf2826 are also present in M. endogenum, and the related Urocystidales *U. primulicola*, and *T. thlaspeos*. In *T. thlaspeos*, an Nlp1, a necrosis inducing protein (THTG_00351; PFAM: PF05630) has been identified as a noncytotoxic effector with preliminary functional validation indicating it contributes to virulence. Nine orthologs are present in *T. frezzii*, five of which are predicted to be secreted, and one of which is predicted to be an effector by EffectorP, ACQY0O_005521, containing the known protein domain, PFAM: PF05630, Necrosis inducing protein (NPP1). *Thecaphora frezzii* has no genes orthologous to TtTue1 (THTG_04687), which encodes a likely effector in *T. thlaspeos*.

Predicted effector-encoding genes often occur near each other in *T. frezzii* (Fig. 1), and are, on average, more likely than BUSCO genes to have another effector-encoding gene within a 100kb window centered on the gene (Table S8). They are also more likely than BUSCO genes to be in repetitive genomic regions (Fig. 1; Table S8), such as near the ends of chromosomes, but are about equally likely to occur near intact transposable elements (TEs), as annotated by EDTA (Table S8). Some effector-encoding genes are organized in clusters, and two have four others within a 100kb window. These clusters occur on two chromosomes, scaffolds 7 and 26. Notably, the three smallest scaffolds, 25, 26, and 27, are mostly comprised of degraded repeat elements, but despite their small size, contain a total of eleven predicted effector-encoding genes. One BUSCO gene is also located on scaffold 27.

### Comparative genomics of smut fungi

Our phylogenomic analysis based on 1112 fungi BUSCO genes resolves the true smut fungi to three clades: one early-diverging lineage containing *Violaceomyces palustris*; the Urocystidales, containing *Urocystis primuliicola*, the two *Thecaphora species*, and *P. flocculosa* Pf-1, sister to *T. thlaspeos* (Fig 2); and the Ustilaginales, containing the other 45 species. Several genera, including *Ustilago* and the anamorphic genus *Pseudozyma*, are not monophyletic, with species assigned to the genera in multiple clades. Smut fungi that have not been observed to be phytopathogenic are phylogenetically diverse and found in all three clades (Fig 2).

Smut genomes vary considerably in size and structural composition. They range in size from 13.0 Mb (*Farysia acheniorum*) to 38.8 Mb (*T. frezzii*: Table S2), in number of predicted genes (5689 in *Melanoaetium* to 10364 in *Ustilago* sp. “UG-2017a”, Table S9), in total intron length of the genome (2.0% in *F. acheniorum* to 8.34% in *V. palustris*; Table S9), and in repeat content (< 0.5% in *Pseudozyma tsukubensis* to 34.5% in *Ustilago* sp.; Table S4). In addition to *T. frezzii*, greater than 10% repeat content occurs in *Anthracocystis setariae* and in 9 species within *Ustilago* (Fig 2; Table S4). All other sequenced Urocystidales have less repeat content (< 5%). Nine of the fifty species have less than 1% repeat content, and 34 have less than 5% repeat content, including *M. maydis*. The least repetitive genomes (< 1% repeat content) have mostly terminal inverted repeats (TIRs) as repeat elements. All genomes with > 5% repeat content have LTRs as the most dominant class of repeats, although the species vary in abundance of LTR superfamilies, as predicted by EDTA. Only *T. frezzii* and *Melanopsichium pennsylvanicum* have notable SINE content, with 3.4% and 1%, respectively.

The 14 species known only as yeasts share genomic characteristics, including few repetitive elements and a significantly higher ratio of secreted proteins predicted as apoplastic effectors to those predicted as cytoplasmic effectors, a trait more typical of saprotrophic fungi (Table 1; Table S10). Lacking significant repeat content, genome size in these species is driven more by genes and introns, while genome size in plant pathogenic species is driven primarily by repetitive elements (Fig 3). The early diverging species also have more introns, inflating the genome sizes.

**Table 1:**
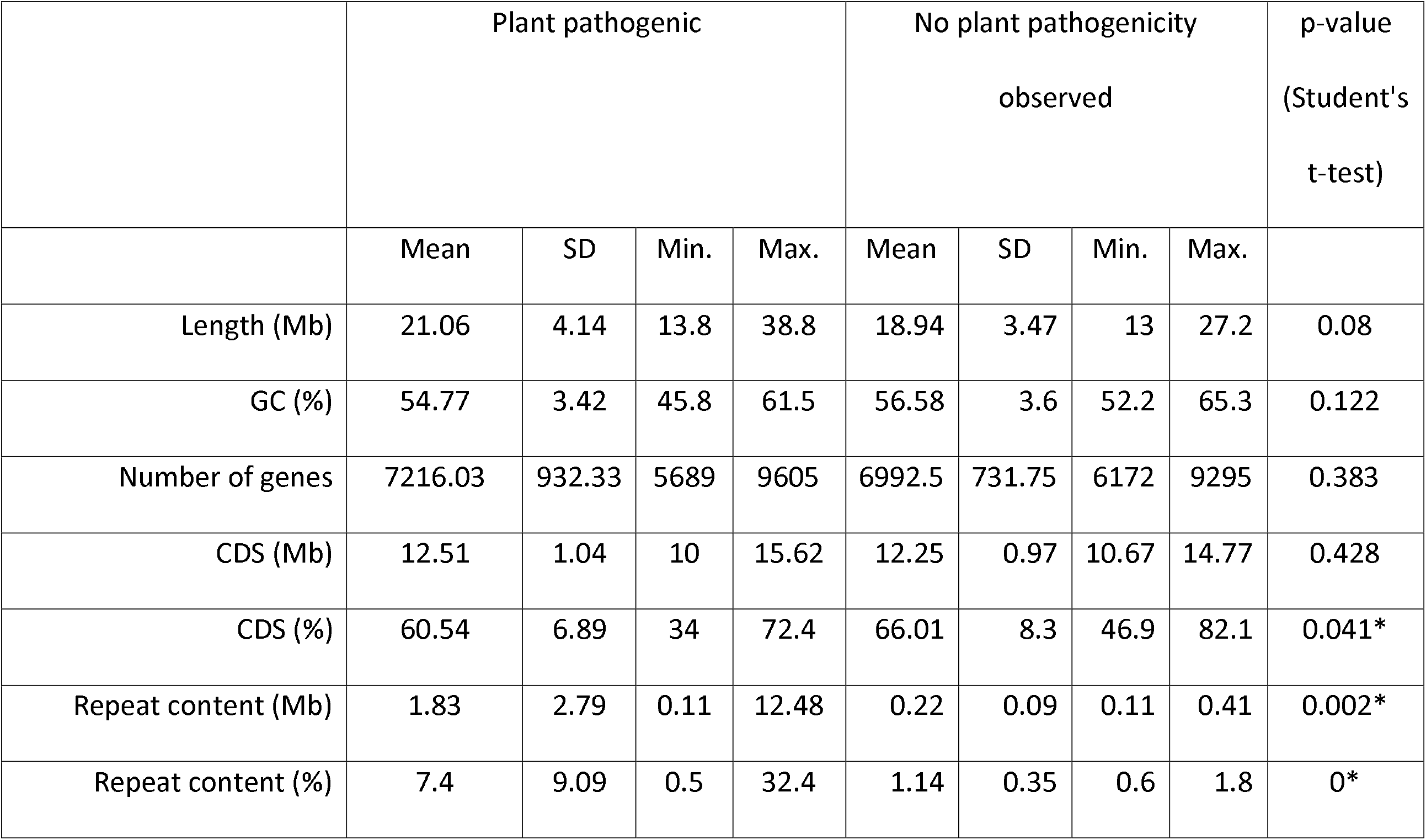
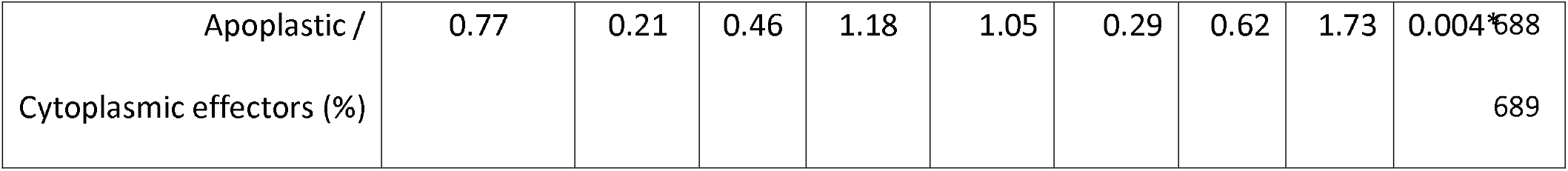
Comparison of genomic features between plant pathogenic true smut fungi and those for which plant pathogenicity has not been observed. Asterisks denote statistically significant differences, based on Student’s t-tests conducted in R. See github.com/ngreatens/Thecaphora_frezzii_genome

Smut fungi share numerous genes confirmed as effectors in the model species *M. maydis*, but many confirmed effectors in *M. maydis* are absent in other species and lineages. Of the core Ustilaginaceae effector orthogroups reported by Schuster et al. (2024), four occurred in all 50 species (Table S6). One of these orthogroups contains an effector known to contribute to virulence in *M. maydis* (UMAG_12205). Eight additional orthogroups were present in 48-49 genomes. Effectors in these eight orthogroups include stp1, stp2, stp3, and pep1, all of which are required for pathogenicity in *M. maydis*. Thirty-two of 57 orthogroups were present in more than 40 genomes. Several other effectors that contribute to pathogenicity in *M. maydis* are not broadly conserved among smut fungi: UMAG_01375 (pit2) is absent in 42 species, and UMAG_12197 (stp4/cce1) is absent in 32 species. The percentage of Ustilaginaceae core orthogroups present negatively correlates with evolutionary distance from *M. maydis* and the Ustilaginaceae (Fig. 2). With the exceptions of the early-diverging *V. palustris* and *Urocystidales*, smut fungi observed only as haploid yeasts retain most Ustilaginaceae core effectors (Table S7).

Conversely, the 150 *T. frezzii* proteins predicted as effectors by EffectorP are within 109 orthogroups (Fig. 2). Fourteen orthogroups are shared among all 50 species, and 32 are present in 48-49 genomes. Sixty-three are unique to the Urocystidales; 58 are unique to *Thecaphora* and *P. flocculosa*; and 39 are unique to *T. frezzii*. Similarly, several families of CAZymes are enriched in the *Thecaphora:* GH28 and PL4 (Table S11).

## Discussion

### The *Thecaphora frezzii* genome

*Thecaphora frezzii* is a serious threat to peanut production in Argentina, a major grower and exporter (Rago et al. 2017). If *T. frezzii* were to spread and establish to other peanut growing regions worldwide, it could cause significant, long-term economic repercussions. To reduce the chance of an incursion by *T. frezzii*, the United States Animal and Plant Health Inspection Service (APHIS) and various other plant protection organizations worldwide have adopted phytosanitary regulations to restrict certain types of peanut imports from Argentina (Arias et al. 2021; Australian Biosecurity Import Conditions 2017; Oilseeds Advisory Committee 2022). As such, it is critical to develop additional resources for phytosanitary efforts. The genomic resources for *T. frezzii* presented here will be a foundational resource for the development of molecular diagnostic tools that will be critical for early detection. Additionally, considerable resources have already been invested in developing peanut cultivars in which resistance to smut is controlled by one or two loci (de Blas et al. 2021; Chamberlin et al. 2024; Massa et al. 2021). While multiple factors affect durability of resistance, single-gene resistance is often not durable (Stuthman et al. 2007). This reference genome will enable researchers to monitor changes in the pathogen population that may threaten host resistance.

In 2023, a 29.3 Mb draft genome of *T. frezzii* was published for the first time (Arias et al. 2023)—an important step toward understanding and managing peanut smut. However, the assembly was fragmented and incomplete. Here, we present a high-quality, chromosome-level assembly of *T. frezzii* with more than 99% of BUSCO genes represented in the assembly. The *T. frezzii* genome is the largest of the true smut fungi genomes sequenced to date and has the most repetitive content (12.5 MB), representing 32.2% of the total genome. Its genome also has a high number of introns relative to other smut fungi, also contributing to its relatively large genome size (Table S9).

Early genome sequencing efforts showed that the genomes of *M. maydis* (Kamper et al. 2006) and the related maize pathogen, *Sporisorium reilianum* (Schirawski et al. 2010), were both small and compact (20 and 18 Mb, respectively), with low proportions of repeat content (2.6 % and 1.8 %). Significant repeat content was first observed in *Ustilago hordei*, which has a larger genome at 26 Mb, but is still relatively compact when compared with other fungi (Laurie et al. 2012). While even the largest known smut genomes are much smaller than those of most obligately biotrophic plant pathogens like the rust fungi and powdery mildews (Kijpornyongpan et al. 2018), recently available genomes of smut fungi vary considerably in size and repeat content (Depotter et al. 2022; Steins et al. 2023). The *T. frezzii* reference genome extends these bounds outward considerably. Taken together, these data suggest there may not be a typical smut genome with regard to these features.

Transposable elements are DNA sequences that move from one location in the genome to another. In some species, they proliferate, inflating genome sizes. Transposable elements, such as long terminal repeat (LTR) retrotransposons, DNA transposons, and others, may significantly affect gene function in fungi, as in other eukaryotic organisms (Muszewska et al. 2017), and changing virulences in plant pathogenic fungi are often associated with TEs. In *T. frezzii*, as in many other fungi, effector encoding genes, such as avirulence genes, are often located in repeat-rich regions, where high mutation rates and epigenetic regulation associated with TEs can contribute to changing virulence profiles (Fouché et al. 2022). In U. hordei, for example, a TE insertion interrupted the promoter of an avirulence gene that encoded an effector recognized by the host, leading to downregulation of the gene and resulting in virulence (Ali et al. 2014). While smut fungi lack the canonical signatures of repeat-induced point (RIP) mutations (Horns et al. 2012), a common means of TE-regulation in the Ascomycota, variability of TE abundance between and within lineages may imply some other mechanism of TE-repression.

*Thecaphora frezzii* encodes orthologs for several core effectors within the Ustilaginaceae, including some that are required for virulence or contribute to virulence in *M. maydis* (Schuster et al. 2024), including pep1, stp1, stp2, stp3, and UMAG_12205. These effectors are widely shared among the true smut fungi, including the early diverging Ustilaginales and the Urocystidales. Like the closely related *T. thlaspeos*, which infects the dicots *Arabis* and *Arabidopsis* (Courville et al. 2019; Frantzeskakis et al. 2017), *T. frezzii* also has a unique array of candidate effectors that do not have orthologs in other smut fungi for which genomes are available, suggesting it employs a unique infection strategy.

Eight of 192 predicted CAZymes in *T. frezzii* are predicted to be secreted effectors localized to the apoplast. Four are within the GH28 family, a family expanded in *Thecaphora* that breaks down galactouronic acid derivatives, structural components of pectin, which is a major part of the cell walls of dicots. Another CAZyme family, PL4, occurs only in *Thecaphora*, as noted by Courville et al. (2019). PL4 family CAZymes break down rhamnogalacturonan, another component of pectin. Two predicted apoplastic effectors are within CE5, a family of cutinases. *In planta transcript* profiling of *T. frezzii* in early stages of infection will help to narrow down the list of candidate effectors and identify those critical for virulence. Identifying effectors that mediate disease outcomes potentially opens a new avenue for managing the disease as they could be useful as targets of plant-mediated RNAi or RNAi-based bio-fungicides.

### Comparative genomics of smut fungi

Species in the true smut fungi have diverse nutritional strategies and ecologies. Some grow only as yeasts, often in the phyllosphere (Inacio et al. 2008), while providing benefits to its host plant (Avis and Belanger. 2002). Some, as saprotrophs, can cause rare diseases in humans (Tap et al. 2016; Telles et al. 2021). Others are useful in industrial fermentation (Jeya et al. 2009). However, most known species are plant pathogens. Due to the relevance of smut fungi to agriculture, biology, medicine, and industry, researchers from diverse fields have published genome assemblies of 50 species at the time of writing. In light of our notably large and repetitive *T. frezzii* genome, we analyzed the genomes of these species to discern patterns relating to genome size, repetitive element content, predicted effectorome, and nutritional strategy, while considering evolutionary history.

Repetitive elements and genome size are correlated with fungal nutritional strategies and ecology, with genomes of saprotrophic fungi generally smaller and less repetitive than those of biotrophic plant pathogens (Raffaele and Kamoun, 2012) and ectomycorrhizal and arbuscular mycorrhizal fungi (Miyauchi et al. 2020). Unlike the rust and powdery mildew fungi, smuts can have both saprotrophic and biotrophic stages of their life cycle. Consistent with saprotrophic fungi generally, smut fungi observed only as haploid yeasts lack significant repeat content (Fig 3). Plant pathogenic smut fungi, in contrast, are variable in genome size and repeat content, with some plant pathogenic species repeat-poor and others repeat-rich (Fig 3). Repeat content, however, is also associated with phylogeny in the true smut fungi, with *Ustilago* containing nine of the 11 species analyzed with greater than ten percent repeat content (Fig. 2). Since phylogenetic relationships may explain plant pathogenicity and repeat content, the relationship between nutritional strategy and genome size and repeat elements within the true smut fungi is not certain, and more sampling of diverse taxa will be needed.

Notably, smut fungi not observed as plant pathogens share one additional trait: a greater proportion of predicted effectors are predicted as localized to the apoplast rather than the cytoplasm, a trait more typical of saprotrophic fungi (Sperschneider and Dodds 2022). This lends evidence to the hypothesis that some smut fungi are adapted to a saprotrophic nutritional strategy. The pathogenicity of saprotrophic smut fungi, however, has recently been brought into question by the discovery of teleomorphs, or the sexual reproductive stage, for several species once believed to be asexual and strictly saprotrophic (Tanaka and Honda, 2017; Tanaka et al. 2019). In addition, many species collected as yeasts and once assigned to anamorphic genera may, based on gene content, retain sexual stages and meiotic ability (Steins et al. 2023). As demonstrated with this study, smut fungi known only as yeasts retain core effectors required for pathogenicity in the Ustilaginaceae. Small, repeat-poor genomes may favor saprotrophic growth in the true smut fungi, even if genes required for meiosis and plant pathogenic ability are retained.

Our phylogenomic analysis is limited by a reliance on publicly available genomic datasets, but our results clearly show that the taxonomy of the true smut fungi requires revision. As others have shown (e.g. Benevenuto et al. 2018), several genera are not monophyletic, with *Ustilago, Anthracocystis, Sporisorium*, and *Pseudozyma* species appearing in multiple clades. A more robust phylogenetic analysis based on whole genome sequences, with representation from type species of genera and underrepresented taxa, will be useful to improve systematics in this group.

### Future work

While 50 high-quality smut genomes are available, a vast majority of research on smut fungi has focused on the model species *M. maydis* and related Ustilaginaceae. Future research into *T. frezzii* will benefit from increased attention to the Urocystidales. High-quality genomes are available for just three other species within this order: *T. thlaspeos, Urocystis primulicola*, and *Pseudozyma flocculosa* isolate Pf-1, which groups within Thecaphora despite being assigned to the teleomorphic genus *Anthracocystis*. Continued use of incorrect generic names or the outdated anamorphic genus *Pseudozyma* obscures differences between unrelated smut fungi, including some of medical and agricultural importance with conceivable impacts on treatment or disease management.

While the infection of peanuts by *T. frezzii* occurs reliably under field conditions, laboratory studies are presently limited by the lack of a reliable inoculation protocol. Studies are ongoing to address these issues, with the goal of infecting peanut plants using dikaryotic cultures of *T. frezzii*. A reproducible inoculation method will enable research into in planta gene expression, studies of host-pathogen interactions, the evolution of host resistance, and functional genetics studies. Numerous smut fungi, including the related *P. flocculosa* (Cheng et al. 2001) and *T. thlaspeos* (Plücker et al. 2021), are amenable to protoplast-mediated transformation. Adaptation of transformation methods for *T. frezzii* will be critical to validate gene function, such as for candidate effector-encoding genes.

Other next steps will include the development of molecular based diagnostic assays and population genetics studies to assess diversity and population dynamics of *T. frezzii* in Argentina. Whole genome sequencing of additional isolates will be useful for examining mating type diversity, and structural and allelic diversity. While much work remains to be done with *T. frezzii*, this high-quality reference genome is a significant step forward in the study and control of peanut smut and in the study of smut fungi more broadly.

## Data Availability Statement

Reads are available under Bioproject PRJNA1250501. The annotated *T. frezzii* genome was deposited at DDBJ/ENA/GenBank under the accession JBPEIW000000000. The version described in this paper is version JBPEIW010000000. Data and scripts used to generate the figures are available at github.com/ngreatens/Thecaphora_frezzii_genome. BRAKER3 genome annotations and orthogroups for the 50 smut fungi are available at 10.5281/zenodo.15677049.

## Acknowledgements

Clint Slocum and Andrew Stone (USDA-ARS), Sol Ailen Bovina (FuEDEI), Ellie Zhao and Lisa Sadzewicz (Institute for Genome Sciences, University of Maryland), and Anna Schultz (Phase Genomics) provided technical assistance. The authors also acknowledge members of the Mars Snacking nut science team for valuable support throughout this project. All opinions expressed in this paper are the authors’ and do not necessarily reflect the policies and views of USDA, DOE, or ORAU/ORISE. Mention of trade names or commercial products in this publication is solely for the purpose of providing specific information and does not imply recommendation or endorsement by the USDA, ORAU/ORISE, or DOE. USDA is an equal opportunity provider, employer, and lender.

