## Supplemental Figures for "A chromosome-level genome assembly of *Thecaphora frezzii*, cause of peanut smut, reveals the largest genome among the true smut fungi"

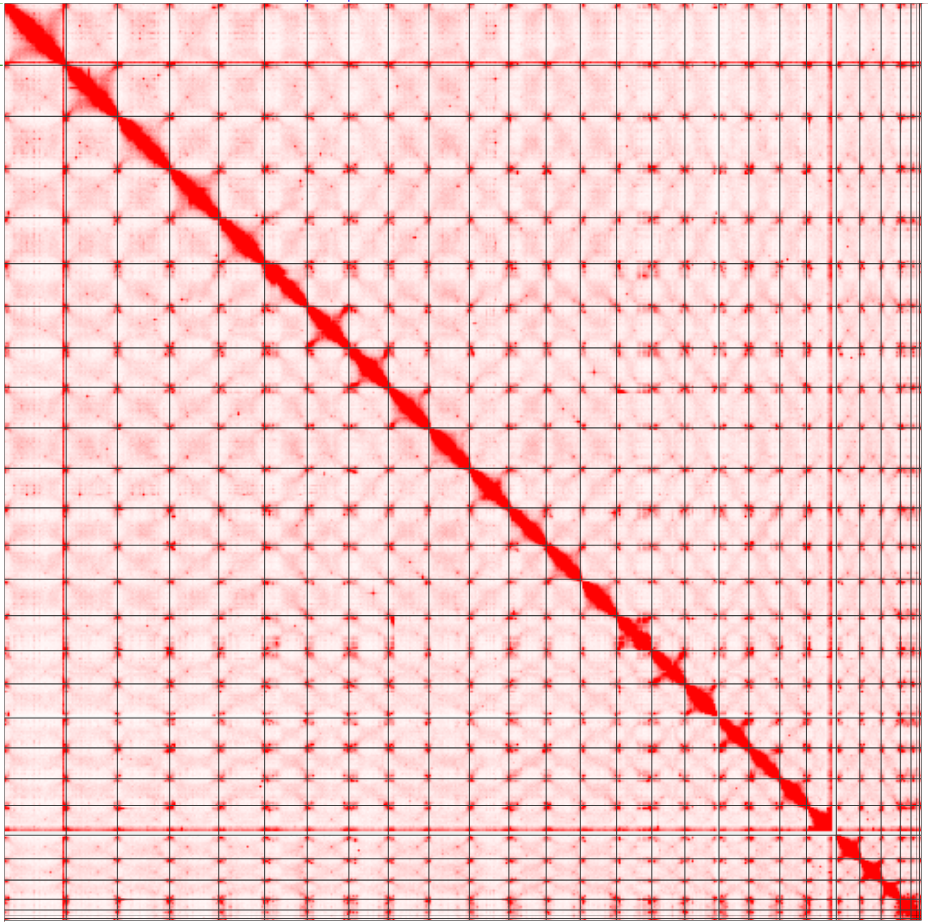

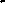

Figure S1. Hi-C contact map for the *Thecaphora frezzii* genome assembly.

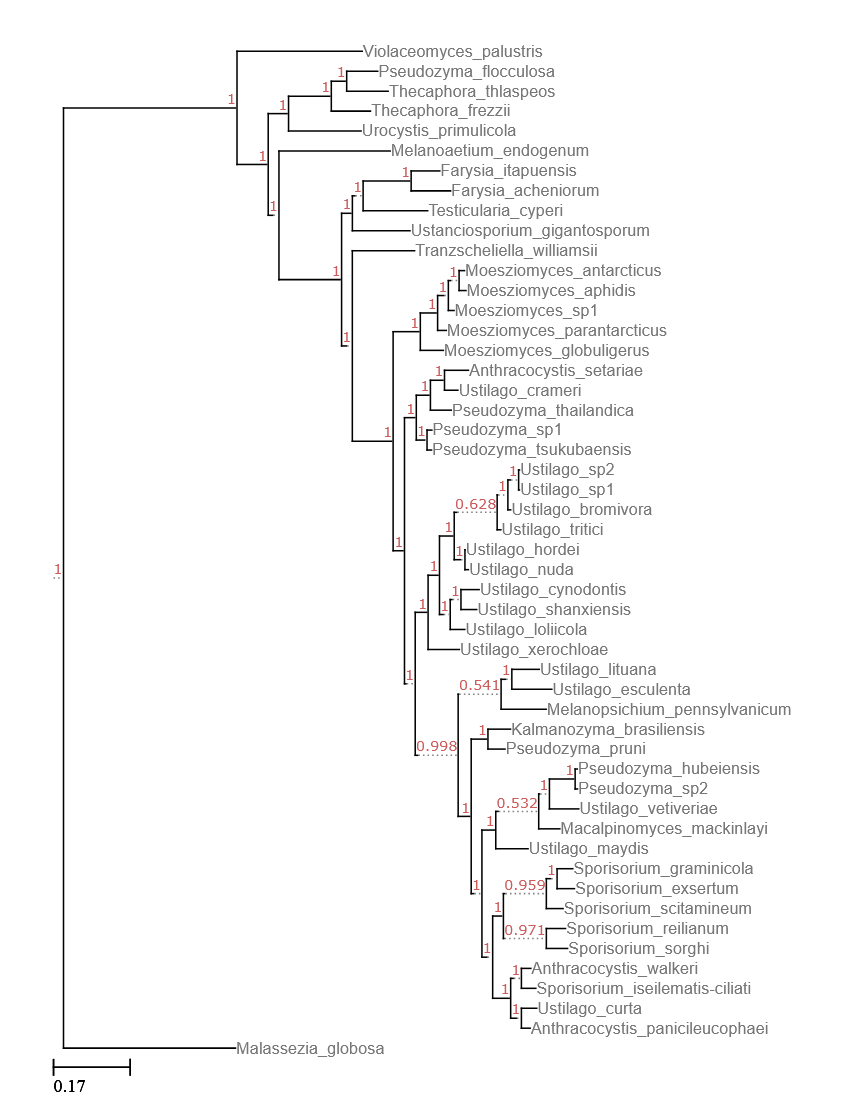

Fig S2. Phylogenetic tree pictured in Fig 1 with branch support values visible.
